# Genome-wide transformation reveals extensive exchange across closely related *Bacillus* species

**DOI:** 10.1101/2023.07.03.547483

**Authors:** Mona Förster, Isabel Rathmann, Melih Yüksel, Jeffrey J. Power, Berenike Maier

## Abstract

Bacterial transformation is an important mode of horizontal gene transfer that helps spread genetic material across species boundaries. Yet, the factors that pose barriers to genome-wide cross-species gene transfer are poorly characterized. Here, we develop a replacement accumulation assay to study the effects of genomic distance on transfer dynamics. Using *Bacillus subtilis* as recipient and various species of the genus *Bacillus* as donors, we find that the rate of orthologous replacement decreases exponentially with the divergence of their core genomes. We reveal that at least 96 % of the *B. subtilis* core genes are accessible to replacement by alleles from *Bacillus spizizenii*. For the more distantly related *Bacillus atrophaeus*, gene replacement events cluster at genomic locations with high sequence identity and preferentially replace ribosomal genes. Orthologous replacement also creates mosaic patterns between donor and recipient genomes, rearranges the genome architecture, and governs gain and loss of accessory genes. We conclude that cross-species gene transfer is dominated by orthologous replacement of core genes which occurs nearly unrestricted between closely related species. At a lower rate, the interplay between the core and accessory genomes gives rise to more complex genome dynamics.

## Introduction

Horizontal gene transfer (HGT) is an important factor in bacterial evolution. It plays a major role in providing non-sexually reproducing organisms with genetic variability. Phylogenetic studies have shown that a surprisingly large fraction of bacterial genomes and gene classes have been affected by HGT over the course of evolution (1-4). However, the genome-wide dynamics of HGT are poorly characterized.

In bacteria, one of the main mechanisms mediating HGT is natural transformation. In this process, cells take up DNA from the surrounding and integrate it into their genome in a process called homologous recombination. *Bacillus subtilis* is among the 80 species known to be competent for transformation (5,6). After external DNA has been taken up into the cytoplasm, the recombination machinery performs a homology search on the genome (7). Microhomologies as short as 8 nt are recognized and further binding of neighboring nucleotides can initiate branch invasion of the genome (8). If branch migration becomes stabilized, a heteroduplex can form as a three-stranded D-loop and be resolved through genome replication (9). Several studies report the formation of mosaic patterns between donor and recipient alleles resulting from this process (10-14). It remains unclear by which mechanism and how frequently these mosaic patterns are formed.

Various processes limit the efficiency of HGT by transformation. At the level of DNA uptake, competence for transformation is tightly controlled in some species including *B. subtilis* (15,16). Once inside the cell, DNA from different strains or species is subject to degradation by restriction modification systems (17,18). Furthermore, the sequence divergence between the donor and recipient alleles has been established as a main barrier to recombination. Multiple studies have approached this dependence by investigating the rate and outcome of transformation for sets of predefined, single genes (19-22). An exponential decrease in the replacement probability was found for different bacterial species (22,23), including *B. subtilis* (19,24) and *Saccharomyces cerevisiae* (20). Likewise, increasing sequence divergence was found to cause decreasing integration lengths (19). Studying the effects of sequences divergence between single orthologous genes is well suited for understanding specific details in the process, yet it falls short of capturing the whole genome transfer dynamics and effects of accessory genes. Genome-wide recombination between different strains and species has been investigated under selective conditions for *B. subtilis* (18,25), *Streptococcus pneumoniae* (13,14) and *Haemophilus influenzae* (26). In these studies, the probability of detecting replacement of a specific gene was dependent on local sequence identity and on its fitness effects. Their disentanglement requires extensive modeling (25). Under minimally selective conditions, the distribution of fitness effects of transformation has been investigated (27), but the underlying genome dynamics were not systematically studied.

Here, we develop a replacement accumulation assay under minimal selection to investigate how the genomic distance between donor and recipient affects genome dynamics. We reveal that the exponential relation between transfer rate and sequence divergence holds for orthologous replacement within the core genomes of *Bacillus* species. Furthermore, we show that the dynamics of the accessory genomes are linked to the rates of orthologous replacement. By pooling the data from a large set of transformation experiments, we investigate cold spots of orthologous replacement. For two closely related *Bacillus* species we demonstrate that nearly the entire core genome is accessible to replacement. Taken together, our work contributes to understanding the barriers to horizontal gene transfer at the level of the entire genome.

## Materials and Methods

### Strains, media and cultivation

We derive the recipient Bs166 strain (*his leu met, amyE::PhscomK(spc), comK::kan, P_comK_gfp*) (25) from *B. subtilis* BD630. In this strain, the master regulator for competence *comK* is deleted and placed under an IPTG-inducible promoter into the *amyE* locus. As donor species, we use *B. spizizenii* 2A9, *Bacillus vallismortis* DSM 11031, *B. atrophaeus* 11A3, *Geobacillus thermoglucosidasius* 2542 wild types obtained from the BGSC and DSMZ. For the competition experiments, a GFP reporter strain (Bs175) was created based on Bs166 (*his leu met, amyE::PhscomK(spc), comK::kan, lacA::PrrnE-gfp (erm)*) (25).

In the replacement accumulation experiment and competition assay, cells were cultivated at 37 °C in competence medium (CM) (28) which was based on Spizizen’s salts (6 g/l KH_2_PO_4_, 14 g/l K_2_HPO_4_, 2 g/l (NH_4_)_2_SO_4_, 1 g/l tri-sodium citrate dihydrate) to which we added 0.5 % D-glucose, 50 µg/ml L-histidine, L-leucine, and L-methionine, 0.02 % casamino acids, 0.1 % yeast extract, and 0.5 mg/ml MgCl·6H_2_O. Liquid cultures were incubated at 37°C and a shaking frequency of 250 rpm. The Infinite M200 plate reader (Tecan, Männedorf, Switzerland) was used to monitor bacterial growth. For picking single clones (single cell bottlenecking), bacteria were grown on solidified plates of lysogeny broth (LB) supplemented with 1.5% agar. Bacteria were inoculated onto the plate and spread by adding small glass beads and shaking for about 15 s.

Genomic DNA needed for the replacement accumulation experiment and for whole genome sequencing was isolated using the DNeasy Blood & Tissue Kit by Qiagen. This yielded DNA fragments with varying lengths, dominated by approximately 30 kbp long fragments.

### Replacement accumulation experiment

In the replacement accumulation experiment, we perform a separate experiment over 20 independent transformation cycles for each of four donor species. For the first cycle of each experiment, we plate Bs166 recipient cells on an LB agar plate, incubate overnight, and pick 8 colonies for the hybrid lineages and 4 as controls. Cells are resuspended in CM in separate wells on a microtiter plate and grown for 2.5 h at 37 °C. We add genomic DNA from the donor species (∼ 1 genome equivalent per recipient cell) together with IPTG to induce competence. Cells take up DNA for 2h and integrate it into their genome, becoming so-called hybrids. The transformation duration mimics the duration of the competent state in wt *B. subtilis* (29). We end the DNA uptake process by diluting the cells and immediately plating on LB agar plates. Plates are incubated overnight and one random colony is picked from each plate to start the next transformation cycle. For the control samples, we perform the same experiment but without adding donor DNA. Every other day, samples are frozen as backups and for later sequencing. 20 transformation cycles are performed consecutively and hybrid strains are whole genome sequenced (Illumina HiSeq, Eurofins) after cycle 10 and 20. We aim at sequencing monoclonal samples to resolve single hybrid lineages. Colonies picked from the plates are not necessarily monoclonal as heteroduplexes formed through transformation are only resolved after plating through cell division (9). For cycle 10 hybrids, detected recombination events pass through an additional filter in the analysis pipeline (look-ahead filter). This filter only retains changes that are also present at a later sequenced time point. For cycle 20 samples, hybrids are plated and picked a second time. The hybrid strains are named according to the supplemented donor DNA. The final set of whole genome sequences comprises 8 independent replicates of each BspizHyb, BvalHyb, GeoHyb, and 7 independent replicates of BatroHyb. Additionally, we analyse 7 representative replicates from the control experiments.

### Detection of genomic replacements through orthologous recombination

For all strains, nucleotide sequences are obtained from NCBI database in fasta-format. For the recipient Bs166 we use NC_000964.3, and for the donors we use NC_014479.1 (*B. spizizenii*), NZ_CP026362.1 (*B. vallismortis*), NC_014639.1 (*B. atrophaeus*), NZ_CP012712.1 (*G. thermoglucosidasius*). Additionally, for the *B. subtilis* recipient, the gene annotation was downloaded in gff3-format. Genes with non-unique locus-tags were removed, leaving 4534 annotated genes. The reference fasta was adjusted to the lab strain by replacing differing SNPs and Indels.

We isolate genomic DNA from all hybrid samples, the recipient and donor species, and obtain whole genome paired-end sequencing reads with a length of 150 bp and an average coverage of 400x. We analyse the data by performing quality control with FastQC (v0.11.7, (30)), trimming with Trimmomatic (v0.36, (31)) and mapping to the appropriate reference with the Burrows-Wheeler alignment tool bwa mem (v0.7.17, (32)). Data is processed with the samtools’ sort function (v1.16, (33)) and variant calling is done with the bcftools’ mpileup and call function (v1.16, (34)). Variants are filtered by a read depth and base quality of at least 50. This pipeline is mostly adapted from the work of Power et al. (25).

With the pipeline, we call variants of the hybrids and donor species against the Bs166 recipient reference. By comparing the single nucleotide changes in the hybrids to the changes between donor and recipient, we detect orthologous recombination as clusters of donor nucleotide alleles (25). These clusters consist of at least 2 donor alleles and end when 5 or more consecutively missing donor alleles are detected. Here, we leave out multi-mapping regions to which reads map inconclusively. We detect these regions beforehand by using Blast (blastn) (35) to align the Bs166 reference genome to itself. We find 10 regions with a total length of about 50 kbp, containing rRNA and tRNA loci. For obtaining the length of the replaced segments, the recombination clusters are extended by half the distance to the next unchanged recipient allele on both sides of the segment. By doing so, we account for the fact that segment integration might initiate at completely identical stretches in which we cannot detect replacement. For each segment, the identity is computed as the fraction of alleles that are unchanged between recipient and donor. For the distribution of sequence identity of the replaced segments, we determine the 95 % confidence intervals of the mean by performing a bootstrap analysis with 10^4^ resampled data sets.

The presence of fully identical stretches prevents us from detecting the exact start and end positions of replacements and thus influences the measurement of segment length and identity. The deviation is strongest for short segments and we thus exclude segments below 100 bp from length and identity analyses.

### Detecting replaced genes and their statistical overrepresentation

For all replaced segments, we use the start and end positions of genes from the annotation of the recipient to detect the affected genes. These gene lists are used to identify overrepresented functional categories with PANTHER GO (36,37). Here, *Bacillus subtilis* is selected as organism and we perform a statistical overrepresentation test with a Fisher’s Exact test and a Bonferroni correction on the PANTHER protein classes.

### Accessory genome and core genome identity

In this study, the genomic distance for each pair of recipient and donor species is quantified by the fraction of shared core genome, the sequence identity within this core genome, and the complimentary accessory genome. These definitions are different from those used in pan-genome analyses, where many genome sequences are compared from strains of the same species (38,39). There, genes shared by all strains make up the core genome, and genes only present in a few strains form the accessory genome (38). Here, we are interested in pairwise comparisons of recipient and donor and, therefore, the accessory genome refers to pairs of species only.

We detect accessory regions that are unique to the recipient on the basis of our pipeline as follows. First, donor sequencing reads are aligned to the Bs166 reference. Then, we detect regions of at least 150 bp length in which the read depth is less than 50. The remaining genomic regions are defined as the core genome in which sequences between donor and recipient are similar and differ only by SNPs. Orthologous recombination can only be detected in the recipient’s core genome. We define accessory genes as those genes that are fully accessory and thus inaccessible to orthologous recombination. Complementary to this, core genes are those that can be affected by recombination and that at least partially contain core genome.

We evaluate the sequence identity between the recipient and donor’s core genome by calling the variants on the whole core genomes and calculating the percentage of similar single nucleotides.

### Detection of deletions, insertions and mixed events

We detect deletions in the hybrids by extracting the coverage with which reads map to the reference with the bedtools2’s function genomeCoverageBed (40). A deletion is detected when the local coverage drops to a coverage of 0 over at least 10 consecutive positions.

Additionally, we are able to detect inserted segments from the donor’s accessory genome in the hybrids. These insertions are facilitated by homologous flanking regions (41). We align the hybrid reads to the donor genome and identify insertions longer than 150 bp through increased coverage in accessory regions.

Deletions of recipient’s accessory parts, insertions of donor’s accessory regions, and replacements can occur alongside each other in the same, mixed integration event. To detect mixed events, the direct proximity of individual events is investigated. For deletions and replacements this is simple, because they are detected with respect to the recipient’s genome. As this is not the case for insertions, we first align the insertion’s homologous flanking regions from the donor to the recipient’s genome. This reveals the most probable entry point for the insertion. Finally, all events are combined, their positioning is analysed and mixed events are identified. In a second approach to connect replacements with insertions and deletions, we align the complete donor genome to the recipient using the Blast algorithm (blastn) and visualize the similarities in so-called dot plots (Fig. S1). Adding the regions of the detected replacement and deletion events on the x-axis (recipient genome), and of insertions on the y-axis (donor genome), we are able to identify mixed events that occurred together due to the positioning and architecture of core and accessory regions on the genomes.

### Detection and analysis of mosaic recombination events and length distributions

We derived the distances between neighboring clusters for the BspizHyb, BvalHyb and BatroHyb samples after 10 and 20 cycles. Here, we determine the transferred lengths by assuming clusters to start and end at the next neighboring missing donor allele (maximum possible extension).

On a logarithmic scale, the distance distribution reveals a bimodal behavior (short-distance mode, long-distance mode). We assume the short-distance mode to represent mosaic replacements that occur together. To dismiss the possibility that these replacement events are instead independent, we investigate a Monte Carlo null model, similarly used by Croucher et al. and others (13,14). For each donor species and time point, we first pool the shortest distances between two events in independent hybrids to ensure independence of transfer distances. From this pool, we collect 10^4^ bootstrap samples that each have the same size as the experimentally detected distance data set, estimate the probability density using a log-normal kernel (to account for the logarithmic x-axis), and count the number of peaks of the probability density. We observe that the distribution of independent distances does not show the bimodal behaviour (Fig. S2a). Then, we perform the analysis to detect the location of the peaks of both distance regimes in the experimental distributions. We draw from the empirical data, perform a kernel density estimation with a log-normal kernel for 10^4^ bootstrap samples, where each sample has the same size as the original data set. For all distributions that are found to be bimodal (for fractions compare Fig. S2a), we collect the position of the peaks and the local minimum between them (Fig. S2b, d). The minimum is used as a threshold to distinguish the short- and long-distance regime. These thresholds are 1340 for BspizHyb, 1940 for BvalHyb, and 10935 for BathHyb. Segments from the short-distance regime are subsequently merged to derive the corrected length distribution of (independent) events.

The length distribution of the recombination events is assumed to follow an exponential probability distribution, described by the single parameter µ, as probability density *p* = µ * *exp*(−µ*x*). Here, we exclude lengths below 100 bp and obtain µ and the error of the fit by fitting the cumulative probability distribution. With µ, we determine the characteristic length, at which the probability is reduced to half of its initial value.

### Obtaining the transfer rates and fitting the exponential relation to the core genome sequence identity

For every transformation hybrid, we fit a linear model to the trajectory of the replaced core genome over time. This is done with a least square fit on a linear model with the intercept set to 0. We test how well the linear model fits the data with the R² quantity that reflects the goodness of fit. For BspizHyb, R² is close to 1 for 4 trajectories and exceeds 0.85 for the remaining samples, except one outlier. For BvalHyb, the value exceeds 0.9 for 6 samples and 0.74 for the remaining two. For BatroHyb, R² cannot be determined, because too many trajectories have values at 0. For each species, we average the slopes of the trajectories. The resulting quantity is called the species transfer rate. Transfer rates *r* are fitted against the sequence divergence (1-identity) to the exponential relation *r*(*x*) = *Aexp*(*B · x*), where the standard deviation serves as weights and A and B are fit parameters. This is done using the scipy.optimize.curve_fit function from python. We obtain the fit parameters, their standard deviation, and the R² value with and without considering the weights.

### Detecting replacement cold spots with pooled data sets

In order to identify genes in Bs166 that have never been replaced by the *B. spizizenii* donor, we draw on additional data sets from hybrids created in different transformation experiments. In total, we use 42 sequenced hybrid organisms. We extend the 8 samples from this study by 12 additional samples created in 20 cycles of the same replacement accumulation experiment. An additional 7 *B. spizizenii* hybrids are included from Power et al. (25). Here, a transformation assay was performed over 21 cycles in which cells were exposed to ultraviolet (UV) radiation on the first day, plated onto agar plates and then transformed with donor DNA on the second day. After this, cells were grown to stationary phase overnight (about 18 h) and plated for single-cell bottlenecking on the next day. The pool is expanded by another 15 hybrids created in 20 cycles of a transformation assay that is similar to the one used in Power et al., but which leaves out the UV radiation step.

For all hybrids, genome replacements are detected as described before. In order to identify gene cold spots, we combine replaced segments from all pooled samples and detect genes that were never replaced. In this analysis, we exclude all recipient genes that contain any accessory or multi-mapping parts, which leaves 3806 core genes. Additionally, we ignore completely identical genes.

### High-throughput competition assay

We measure the relative fitness of the hybrids compared to the Bs166 recipient by performing a recently established high-throughput competition assay (27). This setup allows us to characterize all transformation hybrids in one experimental run and we repeat the experiment on four different days. First, overnight cultures of hybrid strains and the *gfp-*expressing reporter strain (Bs175) are diluted into fresh CM and grown for 2.5 h at 37 °C. After entering the exponential growth phase, the hybrids are mixed in a 1:1 ratio with Bs175 and grown in competition to each other for 4 hours (∼ 14 generations (27)). With a flow cytometer, we measure the fraction of reporter strain *x*_*RS*_and hybrid strain *x*_*i*_before and after competition, at time points *t*_0_ and *t*. Fitness is characterized as selection coefficient *s* with

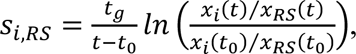

where t_g_ is the generation time of the recipient. By measuring *s* of the recipient Bs166 in each run, we can evaluate the selection coefficient of hybrids with respect to the recipient as *s*_*i,Bs*166_. In total, we measure s for all BspizHyb and BatroHyb strains, and for 7 BvalHyb and 4 GeoHyb strains. Additionally, we perform the assay on 8 control strains that obtained no donor DNA. We use a recently published recipient reference distribution that represents the resolution of the competition assay (27). The data set was obtained by measuring the selection coefficient of 80 ancestor replicates on four different days. The resulting distribution is centered around *s* = 10^−4^ and has a standard deviation of σ = 0.0031.

## Results

### The replacement accumulation experiment is fitness neutral

We sought to find out how genomic DNA from different donor species introduces genetic variations to transforming *B. subtilis*. To this end, we set up a replacement accumulation experiment, using *B. subtilis* (Bs166) as recipient and *B. spizizenii*, *B. vallismortis*, *B. atrophaeus* and *G. thermoglucosidasius* as donors (Fig. 1a). The fraction of the core genome and its sequence identity are used as a measure of the genetic distance between recipient and donor species. In this study, the core and accessory genomes are determined by pair-wise comparison between the recipient genome and each of the donor genomes. Specifically, we determine the fractions of core genomes by aligning donor sequencing reads to the recipient reference and detecting regions where alignment occurs (Materials and Methods). Within these aligning regions, the sequence identity is the fraction of identical base pairs. The remaining accessory genome is unique to each of the species (Fig S1, S4). For the four donor-recipient pairs, the fraction of core genome ranges from 87 % and 86 % for *B. spizizenii* and *B. vallismortis* to 75 % for *B. atrophaeus* and drops to 11 % for *G. thermoglucosidasius* (Fig. 1a). The sequence identities range from 93 % to 80 %. In the following, these distance measures will allow us to detect species-specific differences in the transfer characteristics.

**Fig. 1.**
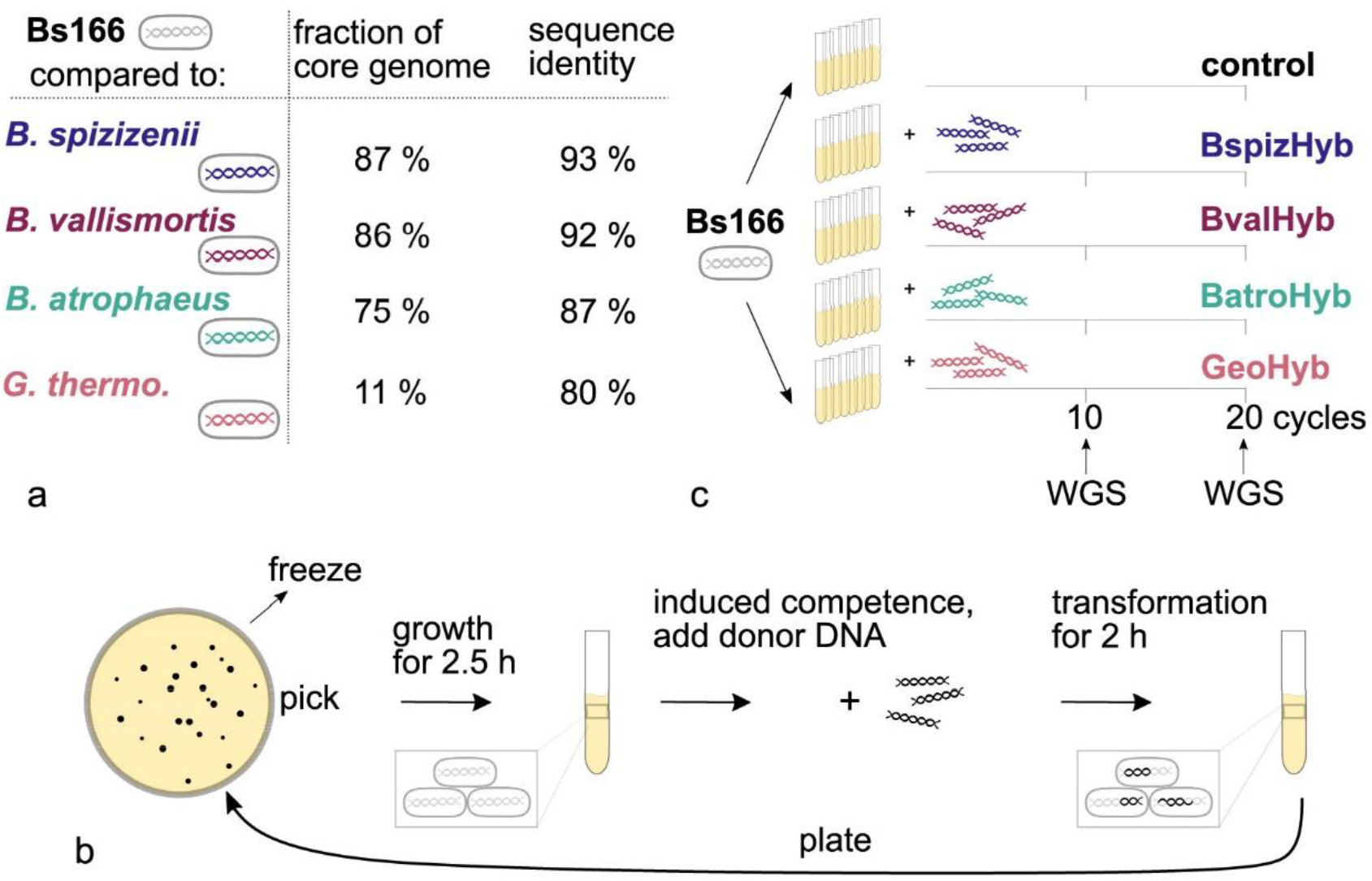
Replacement accumulation experiment performed with four donor species *B. spizizenii*, *B. vallismortis*, *B. atrophaeus* and *G. thermoglucosidasius*. a) Genomic distance between recipient Bs166 and donor species is measured as fraction of core genome and its sequence identity. b) In each cycle of the experiment, one random colony is picked from an agar plate, cells are grown to exponential growth phase, transformed for 2 h with genomic DNA from a donor species, and plated overnight. c) For each donor species, the experiment is performed for 8 replicates in parallel. Additionally, we run a control experiment where cells do not transform with donor DNA. Starting with the recipient Bs166, single transformation cycles are performed consecutively.

The laboratory strain *B. subtilis* 168 develops competence only within a subpopulation comprising up to 10% of the population. Since we are not interested in the 90 % non-transforming recipients, we use a *comK*-inducible strain such that the entire population develops competence upon induction (15). We employ a replacement accumulation experiment (Fig. 1c), because the total fraction of donor sequence in the hybrids is low after a single cycle of transformation (27). For each donor species, we perform the assay with eight replicates that undergo transformation with donor DNA and single-cell bottlenecking in 20 subsequent cycles. A single cycle consists of cell picking, growth to exponential phase, induction of competence, and a period of 2 h in which cells transform with donor DNA (Fig. 1b). Subsequently, the resulting hybrid cells are plated and grown overnight. In parallel, we perform a control experiment in which the recipient is cultivated like the hybrid strains but without adding donor DNA. This replacement accumulation experiment allows us to scale up the fraction of genome being replaced, which is particularly important for distant donor species. The set of hybrids created in the different experiments are called according to the donor organism, BspizHyb, BvalHyb, BatroHyb and GeoHyb (Fig. 1c). After cycles 10 and 20, the hybrids are subjected to whole genome sequencing.

In this experiment, we apply minimal selective pressure as the different hybrids do not compete at any stage. During the transformation step, competent cells are growth arrested (42) and they are plated immediately after the transformation step. From the plates, one single colony is picked randomly and only non-viable variants are lost from the hybrid pool. This experimental design allows us to accumulate unbiased transformation results that represent the general transformation process nearly independently of selection. To verify that the setup indeed is fitness neutral on average, we measured the selection coefficient *s* of hybrids after cycle 10 and 20 with a high-throughput competition assay (Methods) (27). This quantity describes the difference of growth rates per generation between the hybrid strain and the recipient Bs166. For all sets of hybrids, we compare the mean fitness to a recently published recipient reference distribution (n=80) (27). We find that the mean values do not significantly differ from the reference fitness (Fig. 2, Welch’s t-test with a p-value of 5%) at either time point. Inspecting individual hybrid lineages (Fig. S3 a, b), we also find no net increase in fitness over time. Nevertheless, at the level of individual hybrid strains, transformation shows beneficial or deleterious fitness changes in some of the strains. We identify large fitness effects for hybrids by testing against the recipient reference distribution using a Z-test and Bonferroni correction. The number of hybrids showing a strong fitness effect increases from cycle 10 to 20. Overall, transformation causes the fitness distributions to broaden for later time points and higher donor sequence identity, including the BspizHyb and BvalHyb samples (Fig. 2). Taken together, we find no net increase in mean fitness of the hybrid strains, indicating that selection is negligible, and we can study the effects of transformation on genome dynamics independent of selective effects. We note, however, that non-viable hybrids or hybrids that grow too slowly to form a visible colony overnight are ignored in this study.

**Fig. 2.**
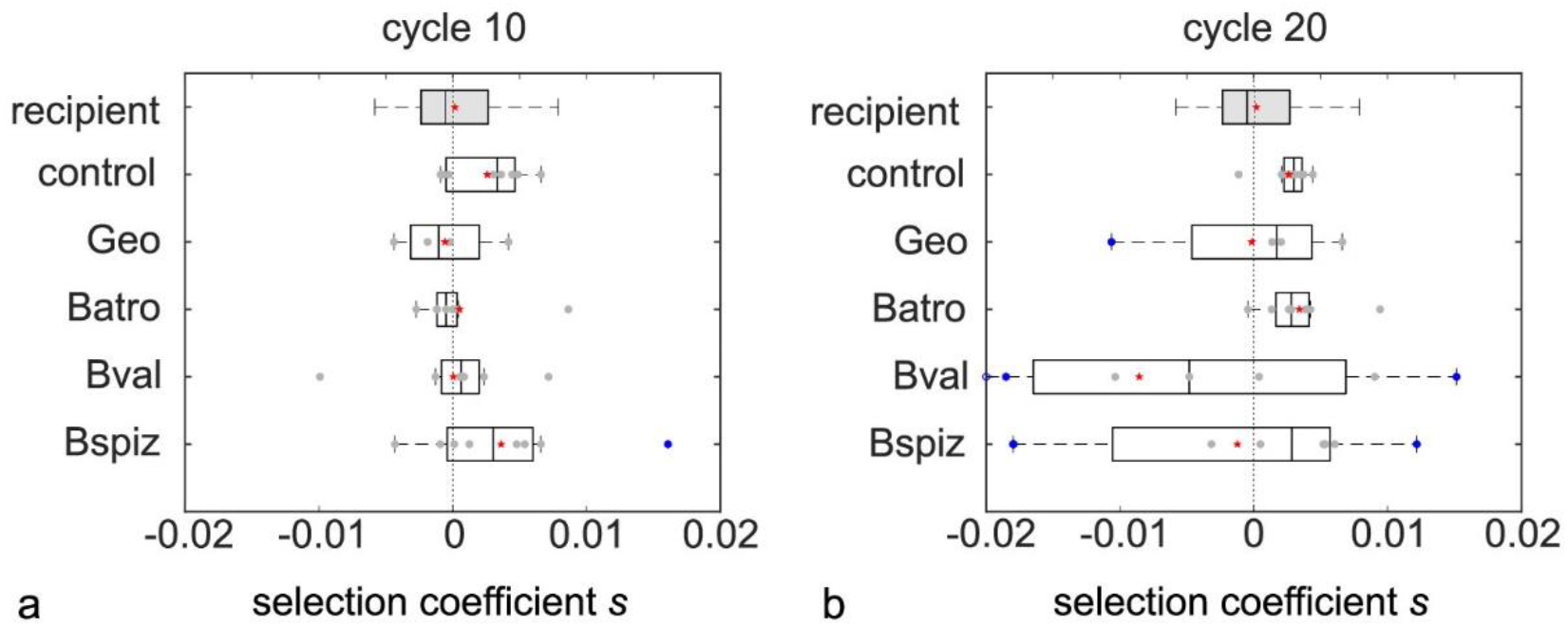
Replacement accumulation does not increase the mean fitness of the hybrid strains. Selection coefficient *s* is shown for hybrids from a) cycle 10 and b) cycle 20. The recipient reference distribution (n=80) (gray box) represents the resolution of the method and is taken from a recent publication (27). For each condition, the box plot with the median (vertical line) and the mean is shown (red star) along with the individual samples (gray circles). The box depicts the first to third quartile of the data. Hybrids with significantly strong effects (blue circle) when compared to the recipient’s distribution are highlighted. For BvalHyb at cycle 20, the extreme negative outlier is depicted on the outer edge of the plot (empty blue circle).

### The replacement probability clusters locally for distantly related *Bacillus* species but not for closely related species

We analyze the outcome of the 20 cycles of the replacement accumulation experiment by detecting three different types of genomic variations (Fig. 3) (25,27). Most frequently, a segment of the recipient’s genome is replaced by the orthologous segment from the donor species. This process is called orthologous replacement and can only be detected within the core genome. Additionally, insertions of segments of the donor’s accessory genome as well as deletions of segments of the recipient’s genome are detected.

**Fig. 3.**
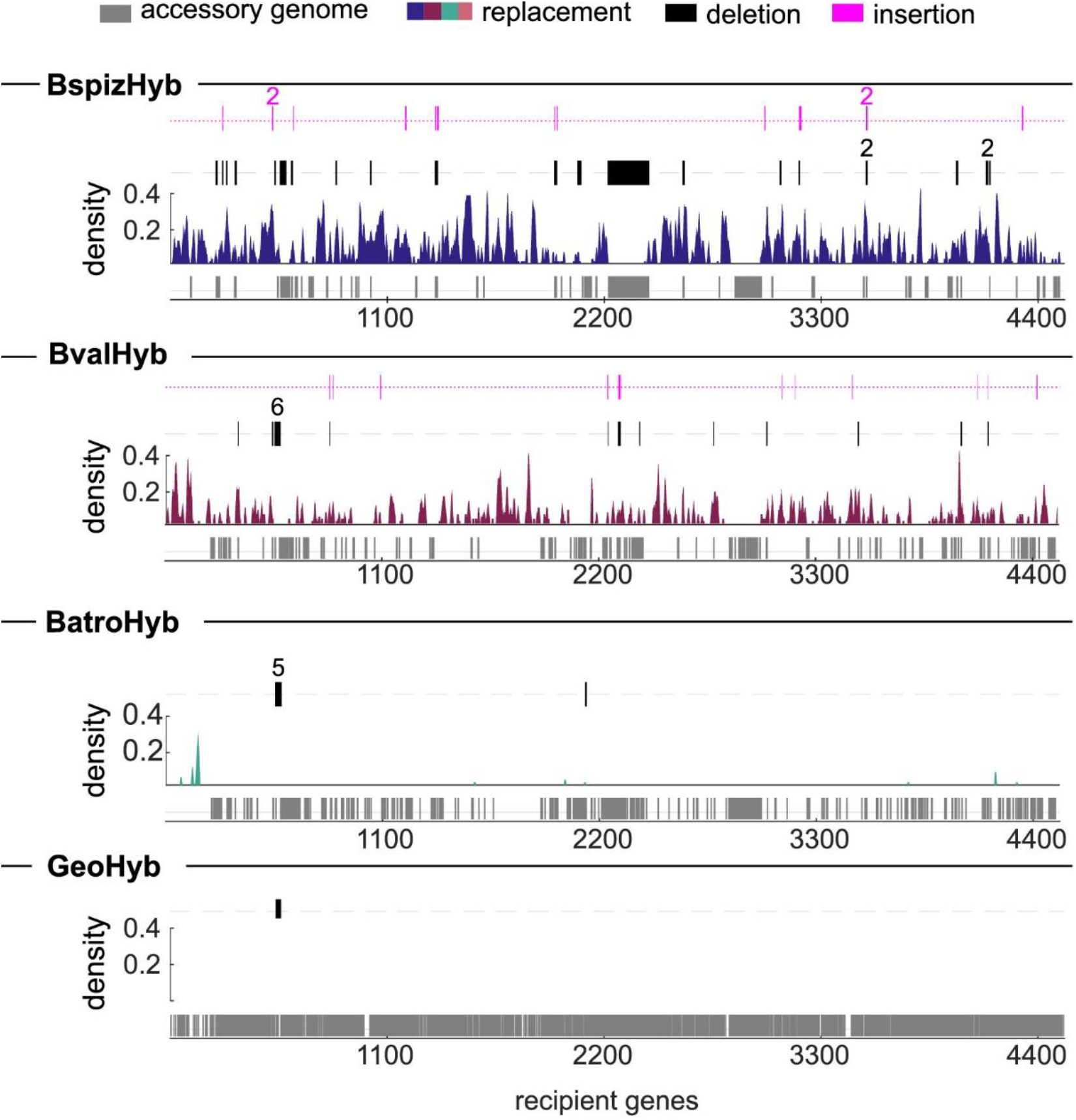
Genomic variations in transformation hybrids depend on the sequence identity between donor and recipient. For all transformation hybrids after 20 cycles, orthologous replacements, deletions, and insertions are detected in the whole genome sequencing data. Across all recipient genes, the recipient’s accessory genes for each donor species are shown in the lowest line (gray bar). The density of genes affected by orthologous replacement is calculated as the average of replaced genes in a sliding window of 10 genes over all replicates. Genes deleted (black bar) or inserted (magenta bar) in at least 1 hybrid strain are shown, whereby numbers indicate multiple hits.

For BspizHyb and BvalHyb, we find that orthologous replacement occurs in abundance along the whole genome, excluding accessory genes (Fig. 3, Fig. S4). Using PANTHER GO, we find no class of replaced genes to be overrepresented. For BatroHyb, very few replacement events are detected. Most of these replacements occur close to the origin of replication, where the sequence identity is particularly high (Fig. S5). Ribosomal genes are overrepresented among the group of replaced genes at a 5 % significance level (using PANTHER GO with Fisher’s Exact test and Bonferroni correction). No orthologous replacement was detected for GeoHyb. The seven control strains that did not receive donor DNA showed no orthologous replacement. Deletions and insertions occur, but their abundance is considerably lower compared to the abundance of orthologous replacement and decreases with increasing distance between donor and recipient (Fig 3). Since most of the insertions and deletions are related to the dynamics of the accessory genomes, we will describe them in detail in a separate paragraph.

In summary, we find that orthologous replacement dominates horizontal gene transfer between different *Bacillus* species. While we observe genome-wide replacements when *B. spizizenii* and *B. vallismorts* are used as donors, for *B. atrophaeus* replacement clusters close to the origin of replication where ribosomal gene are located.

### Segments with a higher sequence identity than the genome-wide average are preferably transferred

We address the question whether the measured sequence identities of replaced segments reflect the average sequence identity between the donor and the recipient’s core genome. To investigate this, we obtain the identity distributions of the orthologously replaced segments after cycle 20 and evaluate the confidence intervals for the mean with a bootstrap analysis (Fig. S6). We then compare the measured identity to the expected core genome sequence identity (Fig. 4). For BspizHyb strains, the replaced segments have an identity of 93.3% [93.1%, 93.4%] and are thus in accordance with the donor’s identity of 93.2%. The numbers in brackets denote the confidence interval of the mean. For BvalHyb, the segment identity of 92.5% [92.2%, 92.7%] is higher than the donor’s identity of 91.7%. Thus, segments with a sequence identity above average are preferentially replaced. This shift between the identity of replaced segments of 92.5% [91.1%, 93.9%] and the core genome identity of 86.5% is even more pronounced for BatroHyb. Examining segments across all samples, we find the minimum detected identity to be around 80 %.

**Fig. 4.**
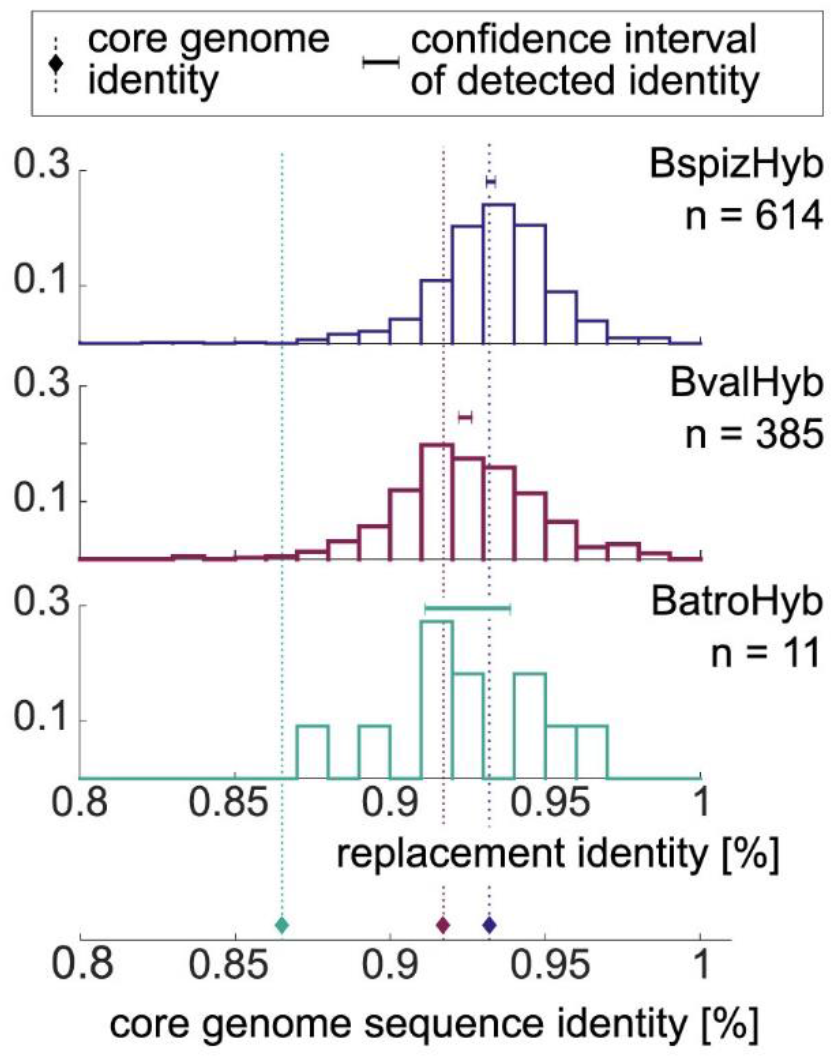

Replaced segments have a higher identity than the average core genome. Normalized identity distributions of transferred segments are shown for the cycle 20 transformation hybrids BspizHyb, BvalHyb, and BatroHyb. Only segments longer than 100 bp are taken into consideration and the sample size *n* differs strongly between the data sets. For each distribution, the 95 % confidence interval for the mean identity is shown (obtained with bootstrap analysis, Methods, Fig. S6). On the bottom, the core genome sequence identity is depicted (diamonds, dotted lines).

We note that we cannot detect recombination taking place in regions with 100 % identity between donor and recipient. However, most completely identical regions are short and replacements detected in our analysis bridge them. Additionally, completely identical regions cause an uncertainty for the localization of recombination start and end sites. As the relative uncertainty is larger for short segments, we exclude those below 100 bp (Methods) in the analysis. Nevertheless, we expect to slightly underestimate the average sequence identity of the replaced segments in Fig. 4.

In conclusion, we observe that the average identity of replaced segments is higher than the core genome sequence identity. This shift increases for donors with higher genomic distances such that, on average, replaced segments from different donors have an identity of around 93 %.

### Transfer lengths are longer for donors with a higher sequence identity

In the next step, we investigate whether the donor species influences the length distribution of transferred segments. When analyzing individual orthologous replacements, we notice some to occur in very close proximity to one another. This is most conspicuous for samples with very few replacements that occur close by. To elucidate this phenomenon, we investigate the distances between neighboring replacements (Fig. 5a). On a logarithmic x-scale, we find a bimodal distribution of transfer distances for BspizHyb and BvalHyb after 20 cycles of recombination (Fig. 5b). The long-distance mode contains more data points than the short-distance mode. Due to the small sample size, the effect is not detectable for BatroHyb. For the two regimes, we assume the large one to contain independently occurring events. We show that the events corresponding to the short-length mode most likely have not occurred independently (Methods, Fig. S2a), indicating that these imports originate from single recombination events that create a mosaic pattern between donor and recipient sequences, as described before (10,11,13). For each donor, (25 - 30) % of all detected replacement events are mosaic events (BspizHyb: 29 %, BvalHyb: 25 %, BatroHyb: 30 %). This suggests that the occurrence of mosaics is independent of donor-recipient identities. Most mosaic events contain 2 replaced segments, and a few have more, ranging up to a maximum of 8 segments in one event.

**Fig. 5.**
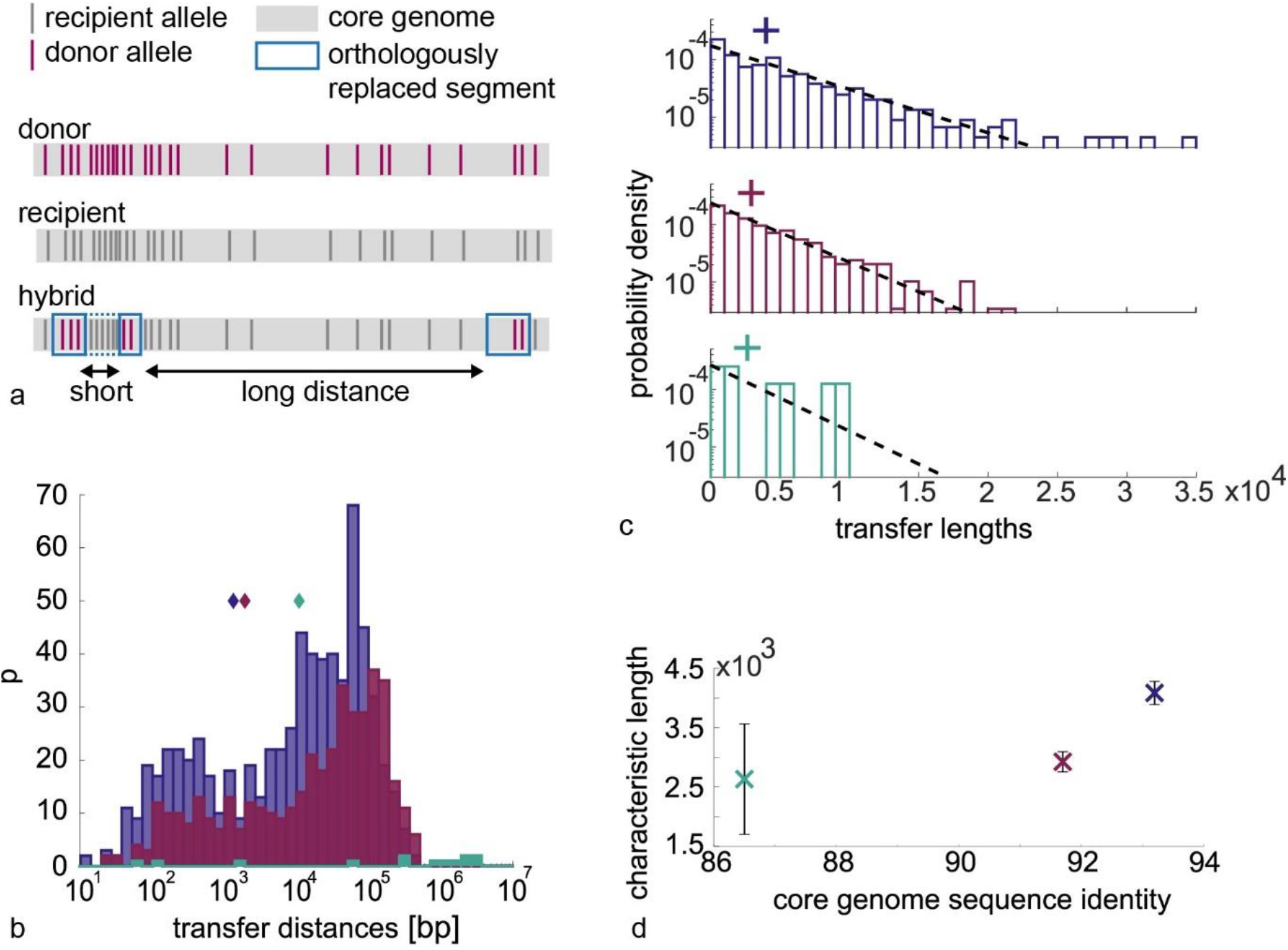
The characteristic transfer length of recombination events increases with increasing core genome sequence identity. a) Scheme of mosaic pattern of orthologous replacement. b) The distribution of distances between replacements, where the cut-off between short and long-distance regime is detected (diamond) using a kernel density estimation (Methods, Fig. S2). c) The length distribution of merged events is fitted with an exponential probability distribution (dashed line) for events longer than 100 bp. The characteristic length is determined (cross) which increases d) with increasing core genome sequence identity (error bars depict the standard deviation). Purple: BspizHyb, magenta: BvalHyb, green: BatroHyb.

Here, we are interested in the length distribution of full recombination events and not in the effects of additional processes that act upon recombination. Thus, we merge the mosaic segments that fall underneath a cut-off value in the distance distribution (Fig. 5b, Fig. S2b, Methods). The length distribution agrees with an exponential decay as a function of segment length (Fig. 5c). The characteristic transfer length increases with increasing donor sequence identity (Fig. 5d). Qualitatively, we find the same relation in the uncorrected length distribution, where mosaic replacements are considered as independent replacement events (Fig. S7). This exponential distribution is expected, if we assume that the length of a segment is determined by the rate of RecA filament elongation and a constant probability of termination (11,43). It is reasonable to assume that the termination rate increases as function of sequence divergence, explaining why the characteristic length of segments decays as a function of divergence as observed here. We also note that the probability density of segments shorter than 100 bp deviates from the exponential relation. It has been suggested earlier that these short segments are integrated by an alternative mechanism (44).

### Gene transfer rate decreases exponentially with core sequence divergence

We address the question how the rate of orthologous replacement (aka transfer rate) depends on core genome divergence. The transfer rate is measured using our time-resolved sequencing data of the replacement accumulation experiments. For each hybrid lineage at cycle 10 and 20 (Fig. 1c), we calculate the fraction of core genome replaced by recombination (excluding GeoHyb, where no transfer was detected). Here, each cycle includes 2h of DNA uptake. The depicted hybrid uptake trajectories (Fig. 6a) reveal an approximately linear increase over time. Even though transfer rates differ strongly, the linear characteristics of the uptake process are preserved between donor species (Methods). We obtain a transfer rate for each trajectory as the slope from a linear fit.

**Fig. 6.**
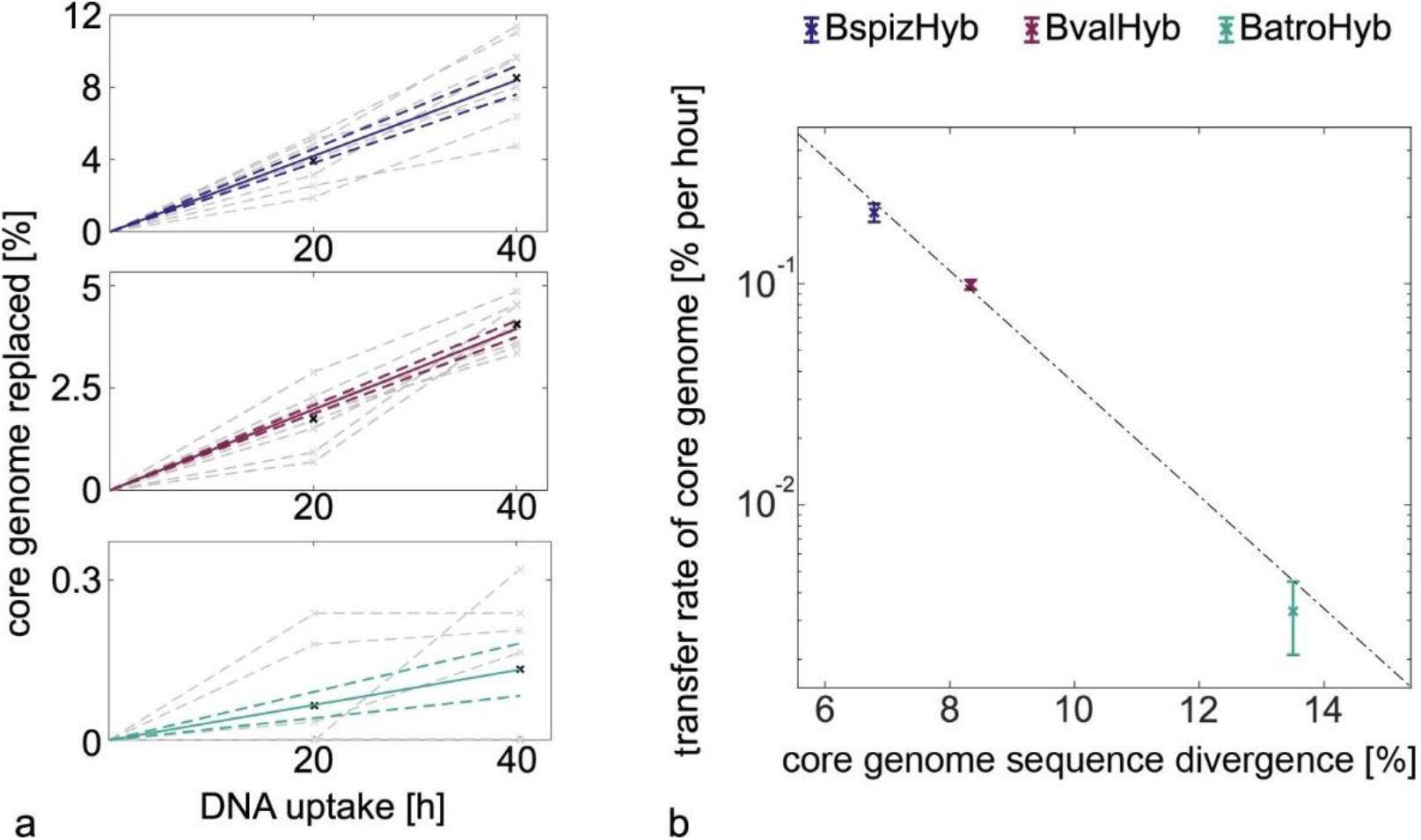
The transfer rate depends exponentially on the average sequence divergence between donor and recipient. a) For different donors in the replacement accumulation experiment, the fraction of core genome replaced increases linearly with hours of DNA uptake. Each hybrid’s trajectory is shown separately (gray line/ cross) together with the mean (black cross). For each hybrid, the data is fitted and the fit mean ± standard error is shown (solid and dashed line). b) Transfer rates decrease exponentially with the sequence divergence of the core genome. Error bars correspond to standard errors. An exponential curve is fitted to the data, yielding an exponential coefficient of −0.59%⁻¹ with a standard deviation of 0.11%⁻¹and an intercept of 12.45%ℎ⁻¹ with a standard deviation of 10.85%ℎ⁻¹(R² = 0.974, weighted R² = 0.999).

The species transfer rate *r* is determined as the mean of the individual rates and the error is evaluated as the standard error. For BspizHyb we find a rate of *r* = (0.210 ± 0.020)%ℎ^−1^. The rate of *r* = (0.1000 ± 0.005)%ℎ^−1^ is slightly lower for BvalHyb, and as low as *r* = (0.003 ± 0.001)%ℎ^−1^for BatroHyb. Taken together, we find an exponential decrease of transfer rate with increasing sequence divergence (Fig. 6b). By fitting this exponential relation (Methods), we detect the exponential decay parameter as −0.59%⁻¹ (standard deviation of 0.11%⁻¹).

In summary, hybrids accumulate orthologous replacements linearly over time and the transfer rate decreases exponentially as a function of average sequence divergence within the core genome.

### At least 96% of the *B. subtilis* core genes are accessible to orthologous replacement by B. spizizenii alleles

We sought to find out whether large portions of the recipient’s core genome are inaccessible to orthologous replacement from a specific donor. To this end, we pool sequencing data from 42 hybrid genomes between *B. subtilis* and *B. spizizenii* that we obtained in different transformation experiments (Methods). All of the strains had undergone 20 or 21 cycles of transformation. On average, 12.5 % of the 3806 full core genes are affected at least partially by orthologous replacement within each strain (Fig. S8a). About half of the strains (20) were transformed under minimal selective conditions as described in Fig. 1. The other strains were grown for about 18 hours after each transformation cycle, allowing for competition between the strains as described in the Methods. We expect that some transfers are selected for or against in this set of strains. Thus, we will only determine an upper limit for the fraction of genes that are not accessible to replacement. Also, the selective pressure is not expected to be strong, because the probability to find genes replaced in a certain number of samples depends only weakly on the selective conditions (Fig. S8b).

Out of 3806 genes that entirely belong to the core genome, only 137 were never affected by orthologous replacement. We will call these genes “putative cold spots” of orthologous replacement in the following. 96.4 % of the genes were at least partially replaced and 91.8 % were fully replaced in at least one out of 42 strains. At the level of base pairs, 94.5 % of base pairs of the core genome were replaced in at least one strain. The putative cold spots are distributed throughout the genome (Fig. 7). We note that replacement cannot be detected if the identity of the replaced segment is 100 %. However, out of 93 completely identical genes, only 3 are detected as cold spots, indicating that this detection limit has no strong influence on the detected cold spots.

**Fig. 7.**
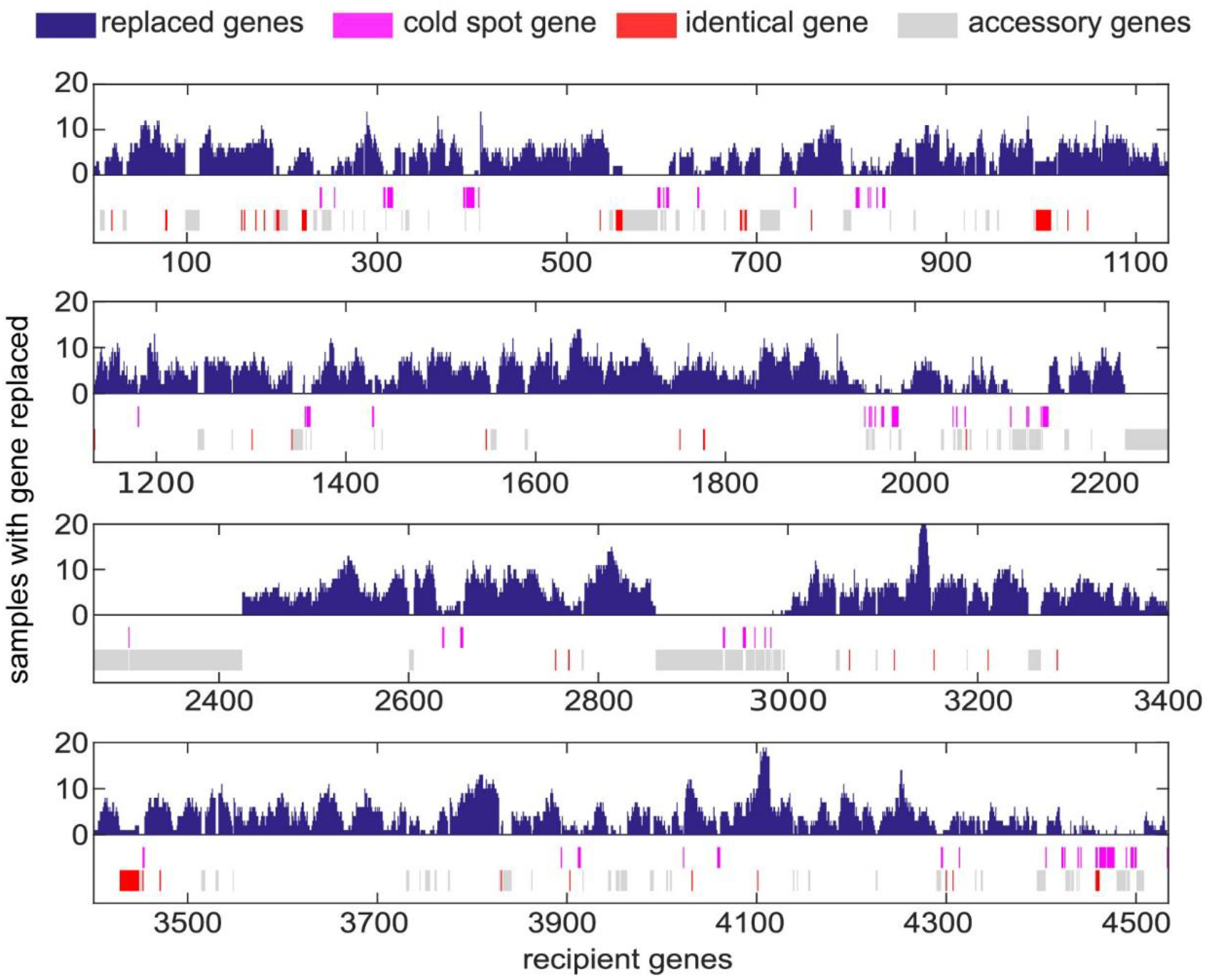
Detection of putative cold spots of recombination with the donor *B. spizizenii* by pooling 42 hybrid strains. Genes that were affected by orthologous replacement in at least 1 out of 42 strains (purple), genes that were affected in 0 out of 42 strains (pink), fully identical genes (red), accessory genes (grey).

Multiple putative cold spots are interspersed between accessory parts of the genome (Fig. 7), indicating that the probability that a recombination event is initiated is affected by the close proximity to non-homologous regions. Therefore, not only the local sequence identity, but also the distribution of the accessory genome affects the replacement probability. Moreover, we find that the distribution of sequence identities of the putative cold spot genes is shifted towards lower identities as compared to the replaced genes (Fig. S9).

We conclude that the largest part of the *B. subtilis* core genome is accessible to recombination with the core genome of *B. spizizenii*.

### The genome architecture influences the transfer of accessory genome parts

Orthologous recombination supports replacement of a core segment by the respective donor part. Additionally, it can also affect the dynamics of the accessory genome depending on its individual architecture (Fig. 8a). Accessory genes of the recipient can be purged through recombination initiated between their flanking regions and donor DNA (Fig. 8b). Similarly, accessory genes of the donor may be inserted into the recipient (Fig. 8c). Finally, accessory recipient genes can be replaced by accessory donor genes and this would be detected as a combined insertion/deletion event (Fig. 8d). In the sequenced hybrids, we find examples for all of the three possibilities (Fig. S1). Their occurrence correlates with the frequency of orthologous replacement (Fig. 8e, f), indicating that they are driven by recombination. Yet they occur at considerably lower frequencies.

**Fig. 8.**
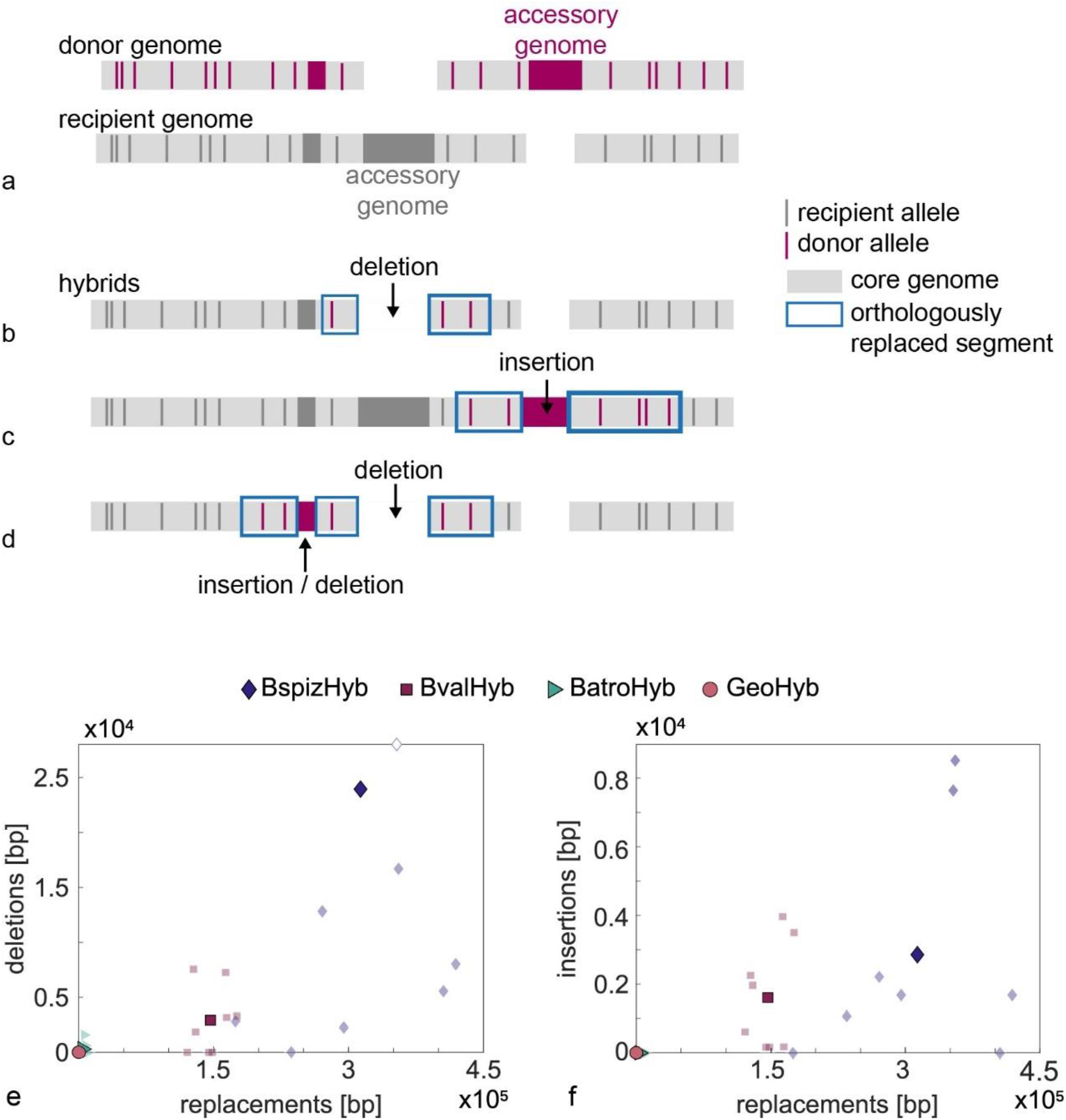
Deletions and insertions occur through orthologous replacements. a - d) Potential rearrangements of accessory genomes through recombination on flanking regions. a) Scheme of donor and recipient genomes. b) Deletion of an accessory segment of the recipient. c) Insertion of an accessory segment of the donor. d) Complex rearrangement in a mixed event including a swap of an accessory segment of the recipient by an accessory segment of the donor and an additional deletion of an accessory segment of the recipient. e, f) For each hybrid after 20 cycles, the number of base pairs deleted (e) and inserted (f) in the transformation hybrids is plotted against the number of base pairs replaced by orthologous replacement. BspizHyb: diamond, BvalHyb: square, BatroHyb: triangle, GeoHyb: circle; light symbols: individual hybrids, dark symbols: average. Mobile element ICEBs1 is excluded from the analysis.

Most deletions purge accessory parts from the genome, as is the case for about 87 % of deleted genes in BspizHyb and 68 % in BvalHyb. For example, in one BspizHyb strain, SPβ prophage comprising about 200 genes on a length of 134 kbp is fully deleted. This happens through recombination that we detect on the much shorter 17 kbp and 7 kbp long flanking regions. Nevertheless, we also find deletions of non-accessory parts when the donor and recipient genomes have different local arrangements, which is the case for both detected deletions found in BatroHyb. Additionally, the mobile element ICEBs1 is lost from the recipient genome under all conditions, but not in all hybrids (Fig. 3, Fig. S4). Even in some GeoHyb samples, where no orthologous replacement is observed, we find occasional loss of ICEBs1. The ICEBs1 element comprises 27 genes and excises itself independently of recombination. Therefore, loss of ICEBs1 is not considered in the analysis of deletions in this study. Excision is known to happen during SOS response after DNA damage due to up-regulation of RecA (45,46), which is also up-regulated during the transformation step in our experiment. Notably, the control samples that did not receive donor DNA did not lose ICEBs1, indicating that the process of transformation induces loss of this mobile element.

We detect all insertions from the accessory genome of the donor to happen alongside orthologous recombination (Fig. 8f) and their frequency is lower compared to the frequency of deletions (Fig. 8e). As the recipient and donor genomes can be rearranged in complicated ways, we find co-occurrence of deletions, insertions and recombination (Methods, Fig. S1) in mixed events (Fig. 8d). For example, in two BspizHyb samples, the stretch containing *comX*, the pheromone involved for example in quorum sensing (47), is replaced by its non-homologous counterpart. These mexed events are rare, yet they reflect the specific architecture of the genomes involved in the recombination process.

We conclude that the arrangement of core and accessory genome influences the dynamics of the accessory genome through transformation. The frequencies of insertions and deletions are low compared to orthologous replacement.

## Discussion

For eukaryotes, mating occurs only between individuals of the same species. This is different for bacteria where genes can be transferred horizontally between species, yet the rates and restrictions of this process are poorly understood at the whole genome level. Here, we employed a replacement accumulation assay for characterizing the effects of sequence divergence on genome-wide transfer between *Bacillus* species. We show that orthologous recombination dominates genome dynamics and that the sequence identity between donor and recipient is crucial for the dynamics of both core and accessory genomes.

For the three *Bacillus* donors studied here, the mean sequence identity of the replaced segments is centered around 93 %. *B. spizizenii* has an average core genome identity of ∼ 93 % with the recipient. Remarkably, nearly the entire core genome of *B. subtilis* is accessible to replacement by *B. spizizenii* core genes. The mean core genome identity of *B. vallismortis* is slightly lower, yet gene transfer occurs across the entire genome. By contrast, for BatroHyb, transfer was observed only within a few genome regions with higher-than-average identity. In these regions, ribosomal genes are overrepresented. It remains difficult to disentangle whether the high sequence identity of these essential genes is a cause or a consequence of enhanced gene transfer rate. These genes may be functional only if their sequence is highly conserved. On the other hand, frequent gene transfer across species may be involved in maintaining the integrity of the essential genes. We found that the rate of replacement decreases exponentially as a function of sequence divergence of the core genomes of the *Bacillus* donors. At the level of single gene replacement, it has been shown that the probability for recombination decreases rapidly as a function of sequence divergence (19,22-24). In those studies, the transformation rate was determined as the number of clones that developed antibiotic resistance due to replacement of a single nucleotide. In the present study, the transfer rate is defined very differently, i.e. as the fraction of recipient core genome replaced by donor genome per time. This rate depends on the local sequence divergence of the replaced segment (comparable to the previous studies), but also on the length distribution of the integrated segments as well as the architecture of the accessory genomes. Therefore, it is remarkable that the exponential relation holds for genome-wide transfer. Given the exponential relation, we can predict the number of nucleotides expected to be replaced when *G. thermoglucosidasius* is used as donor. We estimate a transfer rate of 1.2 · 10^−4^%ℎ^−1^ for GeoHyb, which corresponds to 18 [error 41] transferred bp during the whole of the 40 h experiment. Due to the large error, we cannot determine whether this donor deviates from the exponential behavior and supports the idea of an identity cut-off for transformation as previously proposed (44). Nevertheless, it is interesting to note that we observed no gene transfer across the different genera.

We provide evidence that the dynamics of core and accessory genomes are interconnected. Deletions of the recipient’s accessory genome as well as insertions from the donor’s accessory genome are governed by recombination within flanking regions that belong to the core genome. While these events occur less frequently than orthologous replacement, they generate loss or gain of novel genes and, therefore, are likely to have stronger fitness effects. Vice versa, the architecture of the accessory genomes can also affect the rate of orthologous replacement. We find that multiple putative cold spots of replacement between *B. spizizenii* and *B. subilitis* are located in close proximity to accessory parts of the genome (Fig. 7), indicating that the recombination initiation probability is reduced in these areas.

By comparing the results of the replacement accumulation experiment described in this study with results from our previous study (25), we can assess effects of competition between hybrids on the genome dynamics. That earlier study differed in two important aspects. We let *B. subtilis* transform with genomic DNA of *B. spizizenii*, running 21 cycles. By contrast to the present study, however, during each cycle newly formed hybrid strains were allowed to compete against each other for extended periods of time. As a consequence, the fitter hybrids were selected for. Second, during each cycle, cells were treated with UV light (25). We determined the core genome transfer rate in both studies. Here, we determined a transfer rate of 0.21%ℎ^−1^ for BspizHyb which is lower than the rate found by Power et. al of approximately 0.28%ℎ^−1^. This indicates that high transfer rates were beneficial in the evolution experiment in agreement with the net fitness increase found in Power et al (25). The length distributions of integrated segments were exponential in both studies, but the characteristic length was higher in the earlier study with selection and irradiation than in the replacement accumulation experiment studied here, suggesting that competition has selected for the integration of extended segments. This observation is interesting in the context of disruptive epistasis; with increasing length of the integrated segments, the probability of partially replacing operons decreases. In summary, comparison between two replacement accumulation experiments in the presence and absence of selection, we find a tendency, that selection and / or irradiation enhances the rate of orthologous replacement and the integration of extended segments.

In conclusion, we show that for species with low sequence divergence, nearly all core genes are accessible to gene transfer. The gene replacement rate between *Bacillus* species decreases exponentially with genome divergence between donor and recipient. Orthologous recombination also mediated dynamics of the accessory genomes, however that rate of insertions and deletions of accessory genome was more than a magnitude lower compared to the replacement rate. We propose that cross-species transformation affects bacterial evolution in two ways. First, there is a very frequent transfer between orthologous genes of closely related species. The fitness effects of most of these transfers are fitness-neutral, yet there is potential for beneficial transfer (27). This frequent exchange represents genomic “tinkering”. On the other hand, accessory genes are exchanged at a lower rate, yet we expect them to have larger fitness effects because they enable gain and loss of entire genes or operons.

## Acknowledgements

We thank Viera Kovacova and Leon Haffmans for support with WGS analysis, Lucas Horst and Tobias Bollenbach for help with the liquid handling robot, Gerrit Ansmann for advice on parts of the statistical analysis, and the Maier lab for helpful discussions.

## Funding

This work was supported by the Deutsche Forschungsgemeinschaft through grant CRC 1310 and INST 216/512/1FUGG granted to the Regional Computing Center of the University of Cologne (RRZK) for the High Performance Computing (HPC) system CHEOPS.

## Notes

### Competing Interest Statement

The authors have declared no competing interest.

