## Supplementary Figures for "Genome-wide transformation reveals extensive exchange across closely related *Bacillus* species"

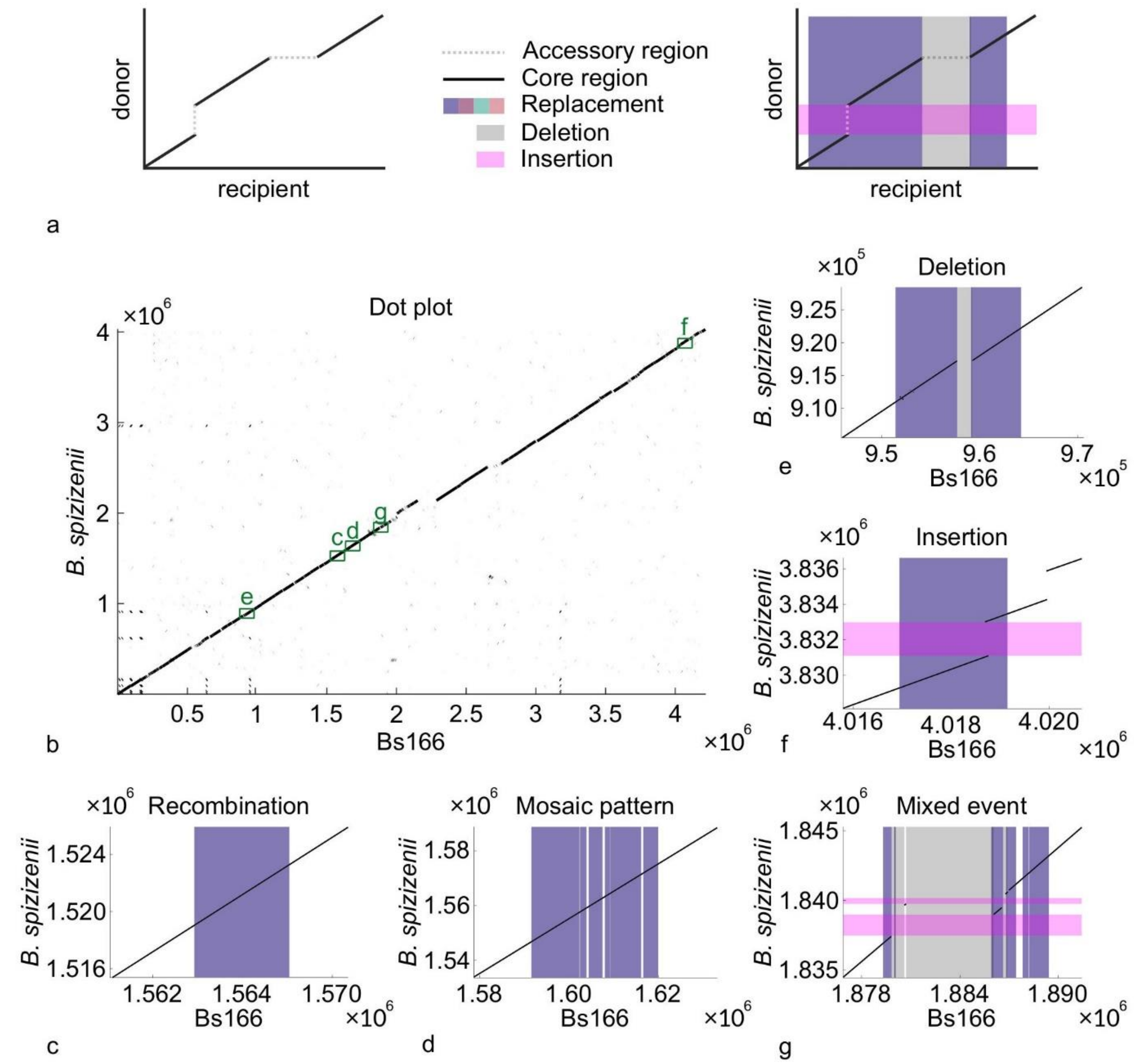


**Fig. S1** Mixed integration events are detected by using dot plots to investigate the differences in genome architecture between strains. Dot plots are created by aligning the donor to the recipient with the blastn algorithm (Methods). a) Scheme of a dot plot in which homologous regions between donor and recipient genome (solid black lines) and accessory regions (gaps, dotted lines) are depicted. Adding detected replacements, deletions, and insertions visualizes the local dependence between events. For *B. spizizenii* and the ancestor Bs166, the whole genome dot plot is shown in b). Most of the homologous regions are present in the same order within both genomes (vertical black line). The green boxes indicate the approximate positions of examples c) – g) that depict genomic changes within the BspizHyb strains. c) contains a simple recombination, d) a recombination with mosaic pattern and, e) and f) a deletion and an insertion both caused by a recombination of homologous flanking regions. In g), we find a mixed event that combines recombination, mosaic, insertion, and deletion.

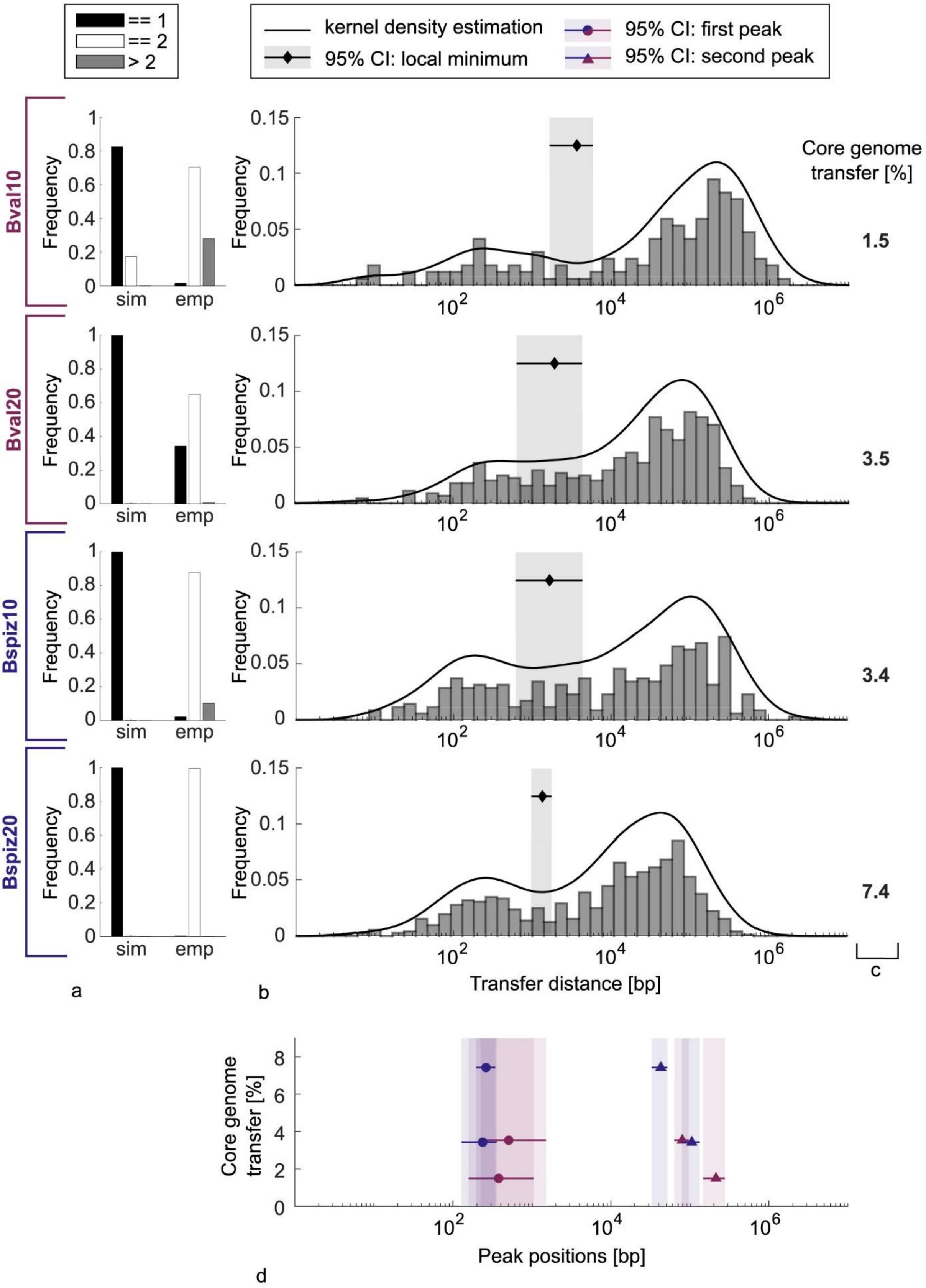


**Fig. S2** The distances between transferred segments reveal a short- and long-distance regime. We assess whether the long-distance regime arises from independently integrated segments whereas the short distances originate from single recombination events that generate a mosaic pattern between donor and recipient alleles. a) By testing a Monte Carlo null model, we ensure that independent transfers do not show bimodal behaviour. In the simulated (sim) independent transfers, most bootstraps only show a single peak (black bar), whereas bimodal behavior (white bar) is most frequent for the empirical data sets (emp). Some bootstraps show more than two peaks (gray bar). b) The empirical data set is represented as histograms and as probability density estimated with a log-normal kernel (shown in arbitrary units). The light gray area around the diamonds depicts the 95 % confidence interval of the local minima. Those minima are used as a cut-off between long- and short-distance regimes. c) The average distance between independent events depends on the fraction of core genome transfer. This is 1.5 % for the BvalHyb after 10 cycles, comparable for BvalHyb after 20 cycles and BspizHyb after 10 cycles with 3.5 % and 3.4 %, respectively, and highest for BspizHyb after 20 cycles with 7.4 %. d) Linking the average core genome transfer of each experiment to its peak positions, we find that the long-distance regime moves to smaller values for increasing transfer whereas the confidence interval of the mosaic peaks overlap for both donors and time points.

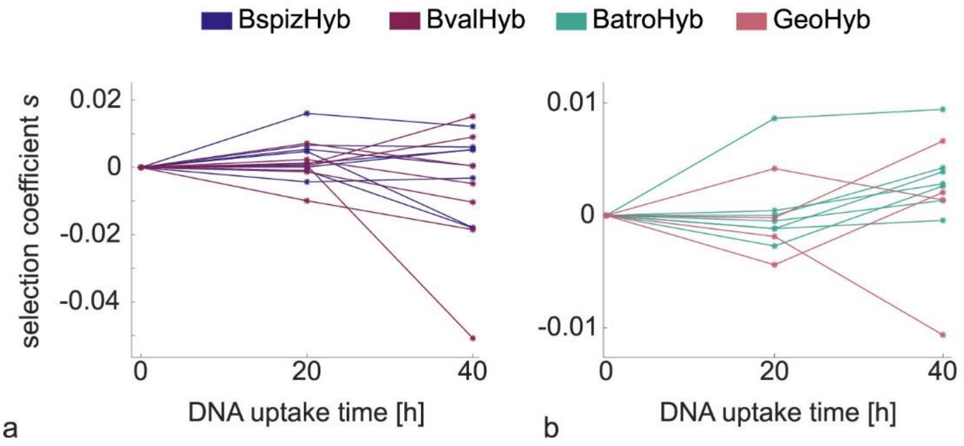


**Fig. S3** Fitness trajectories of transformation hybrids show no net increase. For each hybrid, the fitness is depicted after cycle 10 and 20 as a trajectory. Neither for BspizHyb and BvalHyb a), nor for BatroHyb and GeoHyb b) do we find curves that systematically increase their fitness over time.

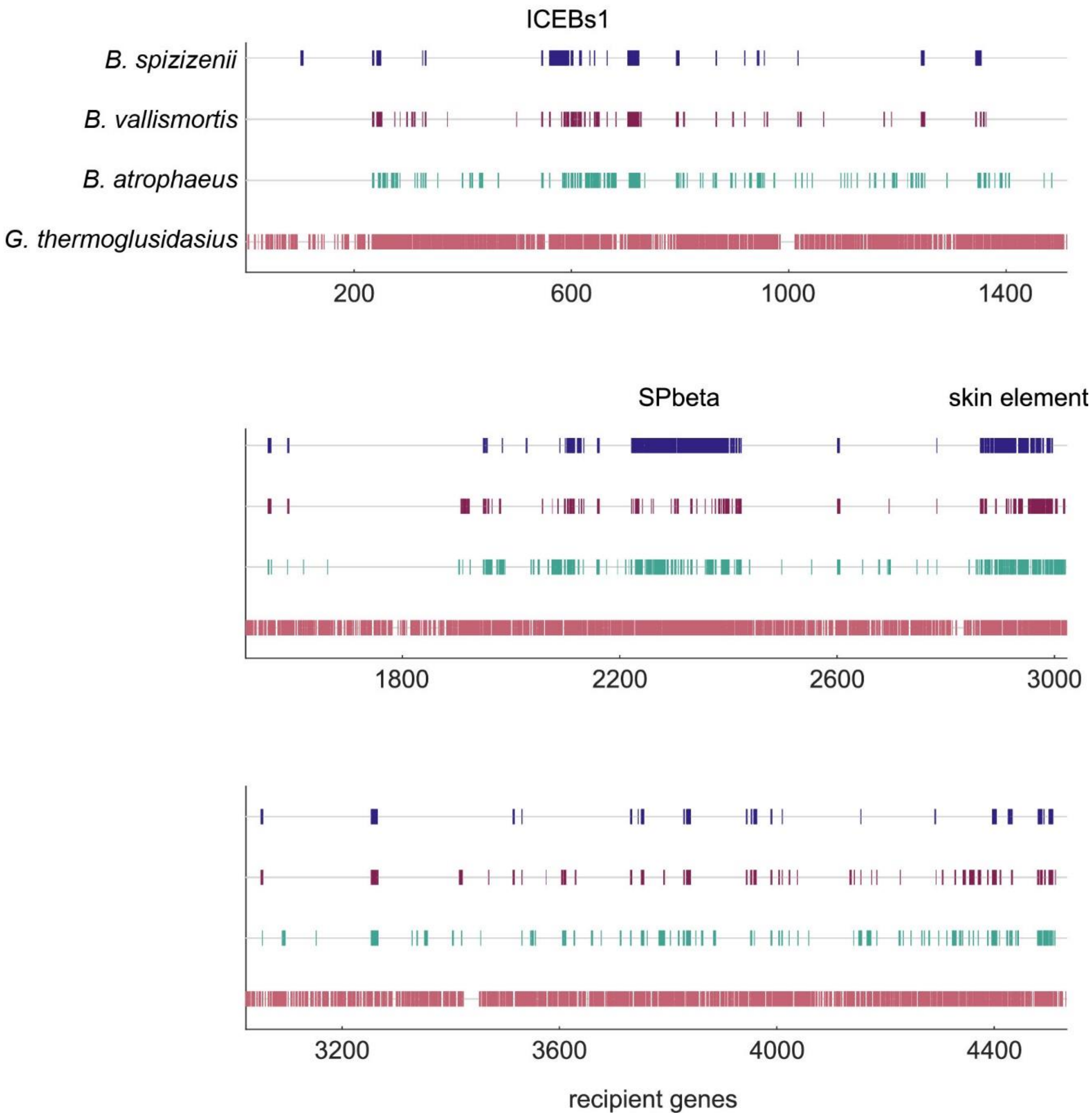


**Fig. S4** Architecture of accessory genomes. The accessory genomes are shown for the Bs166 recipient with respect to the four donor species. Accessory genes are depicted as boxes and the names of important prophage elements and mobile elements are added, according to the annotation (Methods).

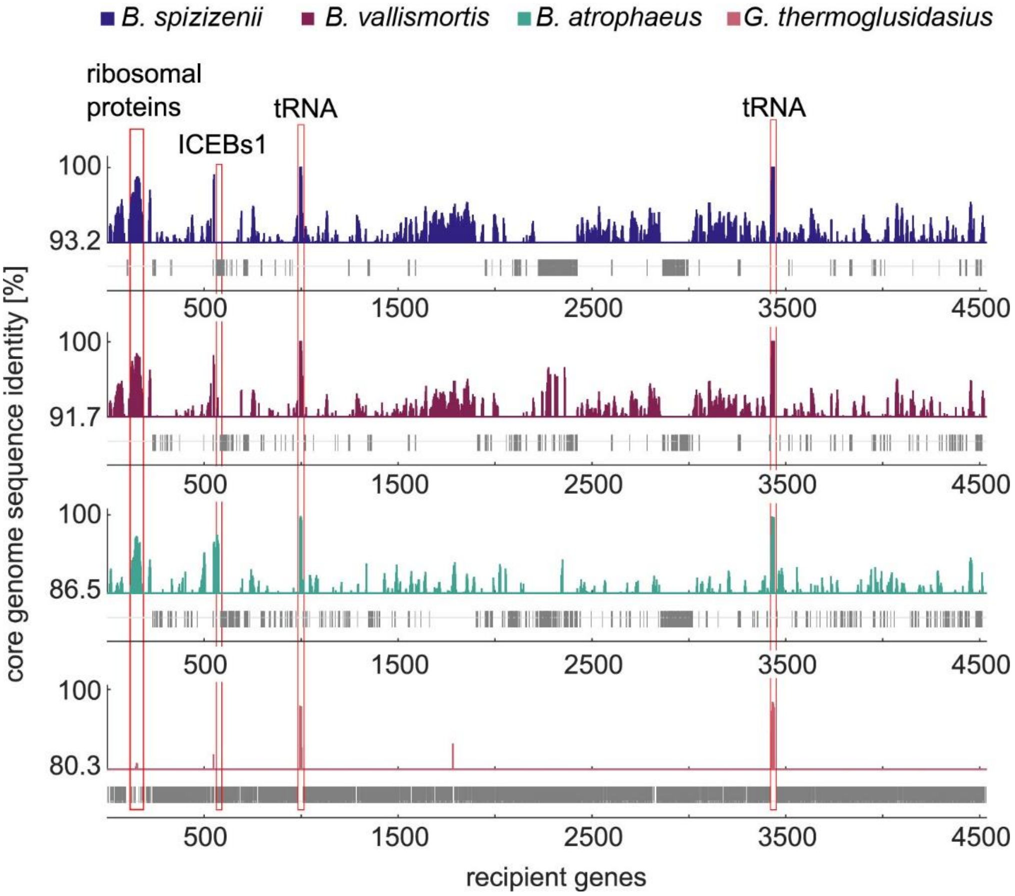


**Fig. S5** The core genome sequence identity is distributed irregularly along the recipient genome. For four donor species with respect to the recipient, the gene-wise sequence identity of the core genome is shown, evaluated as an average in a sliding window of 10 genes. The accessory genes are depicted as gray boxes. The lower limits of the y-axis correspond to the average core genome identities of each species pair. Most prominent clusters of highly identical genes are highlighted.

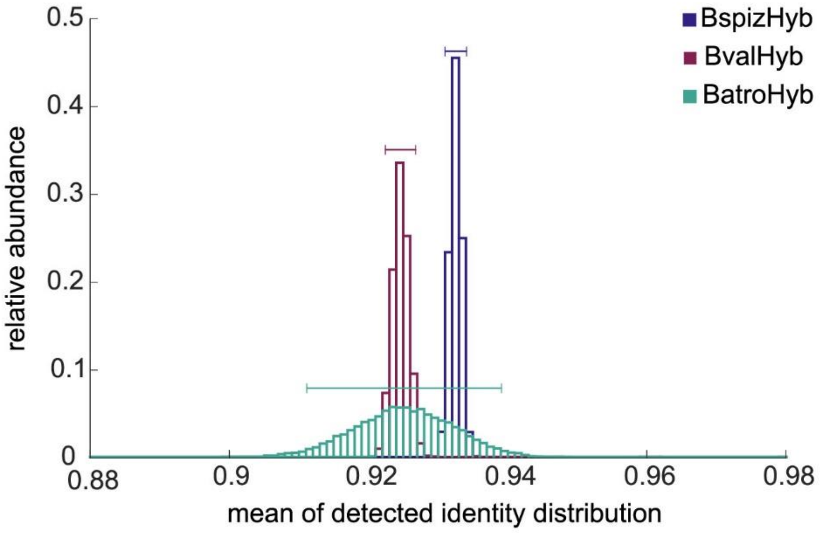


**Fig. S6** A bootstrap analysis for identity distributions of orthologous replacements is performed on the mean of the identity distributions (Fig. 4) for BspizHyb (blue), BvalHyb (red) and BatroHyb (turquoise). Bootstrap samples were drawn $10^4$ times and 95 % confidence intervals are depicted as horizontal lines.

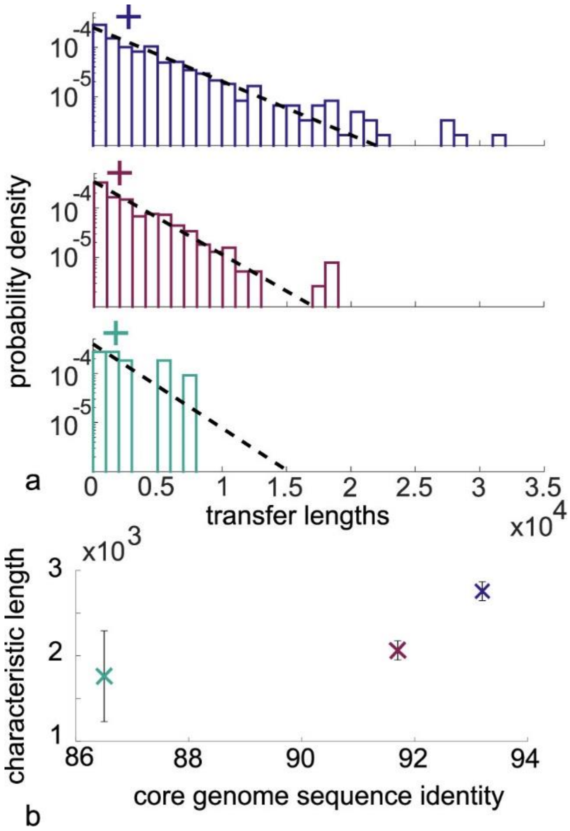


**Fig. S7** Characteristic transfer lengths of replaced segments increase with increasing core genome sequence identity. Here, mosaic segments are not merged. a) The length distribution of segments is fitted with an exponential function (dashed line) for segments longer than 100 bp and the characteristic length is determined (cross). b) This quantity increases with increasing core genome sequence identity (error bars depict the standard deviation). Purple: BspizHyb, magenta: BvalHyb, green: BatroHyb.

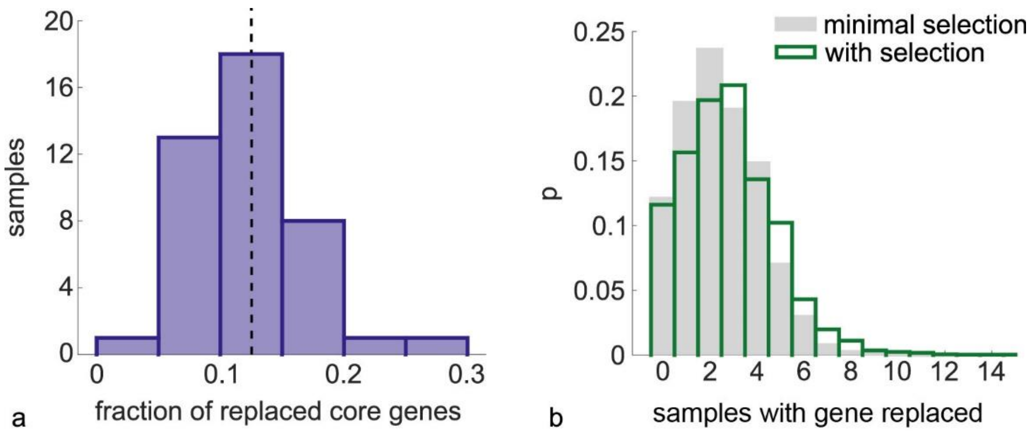


**Fig. S8** Analysis of pooled data from 42 hybrid strains with *B. spizizenii* as donor. a) Distribution of fractions of fully or partially replaced core genes for all hybrids. B) Probability to find genes replaced in a certain number of samples. Hybrids were obtained by single cell bottlenecking under minimal selection (grey) or including competition among different hybrid prior to bottlenecking (green) as described in the Methods.

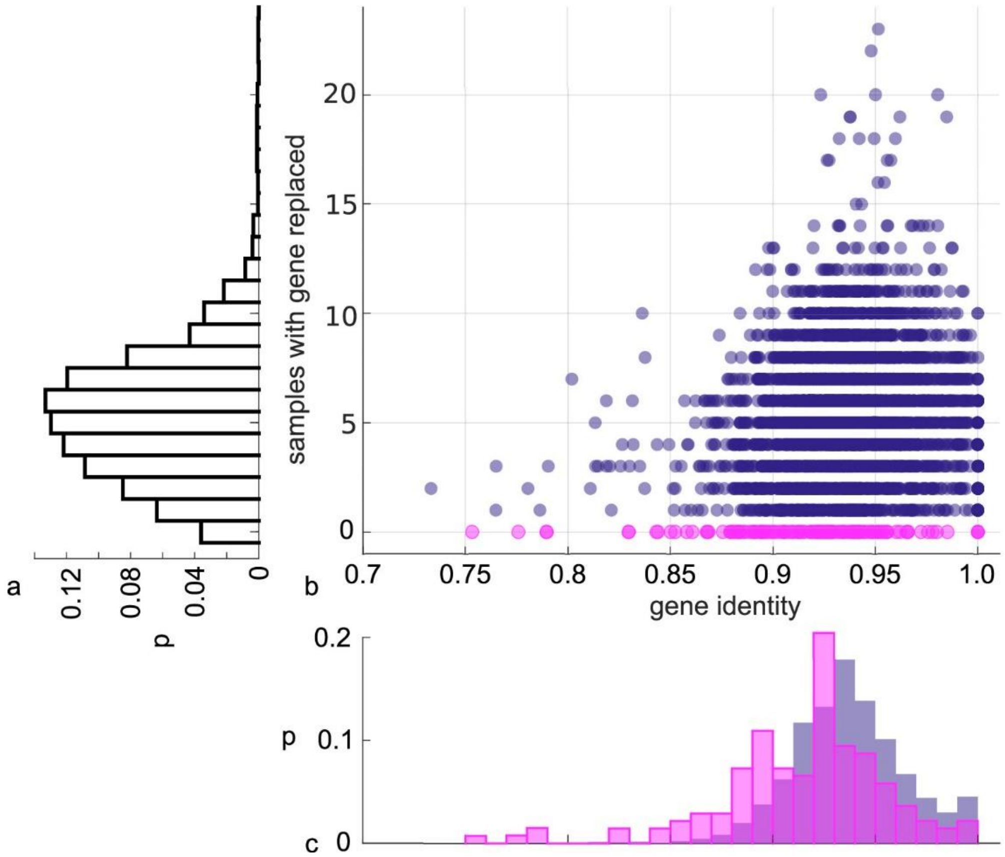


**Fig. S9** Putative cold spots of orthologous replacement have a lower identity than replaced genes. Pooled data from 42 hybrid strains with *B. spizizenii* as donor was analyzed. a) Probability to find genes replaced in a certain number of samples. b) For each core gene, its average sequence identity is plotted against the number of samples it was replaced fully or partially. c) Distribution of gene identity of genes that were replaced in at least one strain (blue) and genes that were not replaced in any of the hybrids (pink). Based on a Welch's t-test, the distributions are different at a significance level of 0.05.
